# Multilevel selection on individual and group social behaviour in the wild

**DOI:** 10.1101/2024.09.16.613329

**Authors:** Conner S. Philson, Julien G. A. Martin, Daniel T. Blumstein

## Abstract

The degree to which phenotypes are shaped by multilevel selection – the theoretical framework proposing natural selection occurs at more than one level of biological organisation – is a classic debate in biology. Though social behaviours are a common theoretical example for multilevel selection, it is unknown if and how multilevel selection acts on sociality in the wild. We studied the relative strength of multilevel selection on both individual behaviour and group social structure, quantified with social networks and 19 years of data from a wild, free-living mammal, the yellow-bellied marmot (*Marmota flaviventer*). Contextual analysis revealed multilevel selection in specific fitness and life history contexts, with selection for group social structure being just as strong, if not stronger, than individual social behaviour. We also found antagonistic multilevel selection within and between levels, potentially explaining why increased sociality is not as beneficial or heritable in this system comparatively to other social taxa. Thus, the evolutionary dynamics of hierarchal or nested biological traits should be assessed at multiple levels simultaneously to tell a more accurate and comprehensive story. Overall, we provide empirical evidence suggesting that multilevel selection acts on social relationships and structures in the wild, and provide direct evidence for a classic, unanswered question in biology.

## 1. Introduction

The evolution of sociality is a central question in biology (Alexander 1974; Wilson 1975; Hinde 1976). Classic explanations for the emergence of social group living focus on the individual fitness costs and benefits incurred via, for example, increased resource competition or predator detection and avoidance (Alexander 1974; Wilson 1975; Hinde 1976). This has helped develop a general theory for the selection pressures of social group living. However, within social groups, individuals vary in their pattern of social interactions with others and this individual variation, in part, drives differences in emergent group social structure (Wilson 1975; Wasserman & Faust 1994; Kurvers et al. 2014; Croft et al. 2016; Kappeler 2019; Cantor et al. 2021). In turn, group structures feedback to influence patterns of individual social interactions (Kappeler 2019; Cantor et al. 2021). Despite these two levels of social organisation being nested, both can be quantified as discrete phenotypic levels of biological organisation (Wilson 1975; Kurvers et al. 2014; Croft et al. 2016; Kappeler 2019; Cantor et al. 2021; Costello et al 2023). These individual and group social phenotypes both have fitness and population dynamic consequences in wild populations (Royle et al. 2012; Kurvers et al. 2014; Croft et al. 2016; Fisher & McAdam 2017; Snyder-Mackler et al. 2020; Philson et al. 2022; Philson & Blumstein 2023a; Philson & Blumstein 2023b) and affect health and human aging (Sapolsky 2004; Snyder-Mackler et al. 2020).

While the drivers and consequences of individual social behaviour and group social structure have been explored independently, if and how these two social phenotypes are under selection relative to each other remains largely unknown. One should avoid to simply do an individual or group gambit simply assuming that the selection estimated on one level is the same throughout all levels. Because the individual and group social phenotypes are nested (like many biological traits), natural selection at one level will likely affect the other (either directly or indirectly), however, selection can also drastically differ among levels (Heisler & Damuth 1987). Selection at one level may increase or inhibit selection at the other, and thus have a stronger impact on the overall evolutionary response (Bijma & Wade 2008). Further, when exploring selection on either level independently, selection on a higher or lower level may be detected at the level being explored, leading to inaccurate or biased evolutionary predictions or false positives. Alternatively, no selection may be detected despite selection occurring at a level not being explored, leading to false negatives and inaccurate evolutionary expectations. Therefore, for nested traits like social behaviour, selection should not be quantified at either level independently but should be quantified simultaneously at all relevant levels to partition selection among the nested levels.

Multilevel selection is a theoretical framework positing natural selection simultaneously occurs at multiple levels of biological organisation, and at levels other than only the gene (Price 1972; Wilson & Sober 1994; Okasha 2005; Wilson & Wilson 2007; Goodnight 2013; Gardner, 2015; Wilson et al. 2023). The theory of multilevel selection has undergone many transformations over the years through healthy debate (Wynne-Edwards 1962; Maynard Smith 1964; Wilson & Sober 1994; Okasha 2005, 2006; Wilson & Wilson 2007; Nowak et al. 2010; Gardner, 2015) and has been suggested as a driving force in the emergence of multicellular organisms (Okasha 2006; Yu et al. 2020), human cultural evolution (Wilson et al. 2023), and the structure of entire ecosystems (Van Vliet & Doebeli 2019; Johnson & Gibson 2021). The modern argument proposes that any phenotypic level, such as the individual or the group, driving individual fitness variation may be under natural selection (Goodnight et al. 1992; Okasha 2006; Gardner 2015; Costello et al. 2023), as seen, for example, in a variety of morphological phenotypes in plants (Weinig et al., 2007) and animals (Eldakar et al., 2010; Fisher et al., 2017; Formica et al. 2021). Individual and group social phenotypes both have documented individual fitness consequences in some birds (Royle et al. 2012), fishes (Solomon-Lane et al. 2015), insects (Costello et al 2023), and mammals (Philson et al. 2022; Philson & Blumstein 2023a; Philson & Blumstein 2023b) and the individual fitness consequences between the two levels don’t always align (Costello et al 2023; Philson & Blumstein 2023b). Thus, multilevel selection may be an important mechanism for the evolution and maintenance of the nested individual social behaviours and group social structures that emerge from group living.

Despite social behaviour being a common theoretical example in making the case for multilevel selection (Wynne-Edwards 1962; Maynard Smith 1964; Wilson & Sober 1994; Okasha 2005, 2006; Wilson & Wilson 2007; Nowak et al. 2010; Gardner, 2015; Wilson et al. 2023), prior work has not explored the presence and strength of multilevel selection for social behaviours in wild, free-living populations. Thus, the decades old debate of individual versus group selection for social behaviours (Leigh 2010), and more recently multilevel selection (Nowak et al. 2010; Goodnight 2013), remains largely a theoretical, not empirical, debate (Okasha 2006). If multilevel selection acts on sociality in the wild is an open question in evolutionary biology that has the potential to inform our understanding of the evolutionary origins of social relationships and structure that underpin social lives across species.

To address if multilevel selection acts on sociality in the wild, we used 19 years of continuous social, fitness, and life history data on a well-studied, wild, free-living population of yellow-bellied marmots (*Marmota flaviventer*). A harem polygynous rodent, yellow-bellied marmots live in colonies and socially interact in summer before hibernating over winter. The evolutionary origins of yellow-bellied marmot group living may be attributable to predator avoidance and/or reflect limited opportunities to disperse to establish new colonies (Armitage 2014). Marmot social groups are comprised of mostly kin, with occasional non-kin immigration (most often a yearling or adult male; Armitage 2014). Individual social behaviour has a genetic basis (Lea et al. 2010) and individual behaviour and group structure are environmentally mediated (Blumstein et al. 2006; Lea et al. 2010; Armitage 2014; Maldonado-Chaparro et al. 2015; Pfau et al. 2023). Individuals also experience fitness consequences based on both their individual and group social phenotypes (Blumstein et al. 2009; Wey & Blumstein 2012; Yang et al. 2017; Blumstein et al. 2018; Montero et al. 2020; Philson et al. 2022; Philson & Blumstein 2023a; Philson & Blumstein 2023b). Namely, more social individuals have increased summer survival due to antipredator benefits (Montero et al. 2020) and are more philopatric potentially due to the costs of dispersal (Blumstein et al. 2009) but have decreased hibernation survival (Yang et al. 2016), reproductive success (Wey & Blumstein 2012), and longevity (Blumstein et al. 2018) potentially due to the time and energetic costs of social interactions. Individuals residing in more socially connected groups also experience decreased individual reproductive success (Philson & Blumstein 2023a), again due to energetic costs, but increased individual winter survival (Philson and Blumstein 2023b) due to potential social hibernation and thermoregulatory benefits. Increased sociality being largely associated with individual fitness costs is largely unique for social mammals (Snyder-Mackler et al. 2020), but nevertheless shows that sociality has consequences in this species.

While exploring nested levels of organisation independently may lead to inaccurate estimation of selection, these studies exploring the levels of sociality independently provide a strong background to ask informed questions about the importance of multilevel selection for social traits. Thus, with identified genetic and environmental drivers of social phenotypic variance and varying fitness consequences for individuals (when explored independently), this system provides an ideal opportunity to estimate multilevel selection for individual and group social phenotypes in the wild to more accurately partition selection among the nested levels of organisation and ultimately obtain a better understanding of the evolution of sociality.

To quantify the social phenotype, we used affiliative social interactions from 172 social groups comprised of 723 unique individuals across 19 years to construct social networks (Wasserman & Faust 1994; Kurvers et al. 2014; Croft et al. 2016) and calculate a pair of analogous individual and group social network measures for four core social traits (Table 1; Costello et al. 2023). Each measure is commonly used across studies in human and non-human animal social network studies (Wasserman & Faust 1994; Kurvers et al. 2014; Croft et al. 2016). The analogous pairs are not strictly required to assess multilevel selection, but because multilevel selection could occur between the social traits, this approach allows clear evaluation of the relative importance of both individual- and group-level selection for specific social traits. To evaluate the distinct contributions of individual social behaviour and group social structure to four annual individual fitness correlates (summer survival, hibernation survival, if a female weaned offspring, and how many offspring a mother had), we used a used contextual analysis which uses a partial regression to partition selection among levels (Lande & Arnold 1983; Goodnight et al. 1992; Costello et al. 2023). Despite the inherent non-independence of the individual and group social phenotypes, contextual analysis partitions among the two phenotypic levels relative to each other.

**Table 1.** The four social traits and corresponding analogous pair of individual and group-level social network measures used to quantify the individual and group social phenotypes. ^*\**^Inverse values of cut points and average path lengths were used so that increase in any of the 8 variables reflected an increase in sociality.

| Social Trait | Individual Social Trait | Group Social Trait |
| --- | --- | --- |
| Connectivity | **Degree:** Number of social relationships | **Density:** Proportion of possible social relationships that are observed |
| Closeness | **Closeness:** Number of social links to access all other individuals in the group | **Inverse Average Path Length<sup>\*</sup>:** Mean social distance between all individuals in the group |
| Breakability | **Embeddedness:** Connectedness in their cluster and group | **Inverse Cut Points<sup>\*</sup>:** number of social relationships that if broken/lost result in two or more separate social groups |
| Clustering | **Clustering Coefficient:** Proportion of an individual's direct social partners who also interact (i.e., local transitivity) | **Transitivity:** Proportion of connected triads actualised |

Because individual social behaviour and group social structure are nested, if we find selection was only observed at one phenotypic level, we could interpret that level as under direct selection while interpreting the other level as under indirect selection (Goodnight et al. 1992). This would not support the presence of direct multilevel selection for social behaviour. If direct selection was observed for both social phenotypes, even across different social traits or contexts, we would have suggestive evidence of multilevel selection for sociality in the wild. Because when exploring the individual and group levels independently, more social individual and group social phenotypes are mostly costly for individual fitness in this system (Blumstein et al. 2009; Wey & Blumstein 2012; Yang et al. 2017; Montero et al. 2020; Philson et al. 2022; Philson & Blumstein 2023a; Philson & Blumstein 2023b), we predicted negative selection (i.e., selection for less sociality) for the connectivity, closeness, and clustering social traits at both levels. We predicted positive selection (i.e., selection for being more social) for breakability traits at both phenotypic levels given more socially embedded individuals and less breakable groups (into two or more social groups) have some individual fitness benefits (philopatry: Blumstein et al. 2009; hibernation survival: Philson & Blumstein 2023b). Lastly, we predicted the strength of selection for individual social behaviour would be stronger than selection for group social structure given the former has more known fitness consequences in this system (when explored independently) and because of the common critique of multilevel selection that the strength of selection for individuals is stronger than for groups (Maynard Smith 1964; Williams 1966).

## 2. Materials and Methods

### (a) Study System

We used a 19-year dataset (2003-2021) on wild, free-living yellow-bellied marmots (*Marmota flaviventer*) studied at and around the Rocky Mountain Biological Laboratory in Colorado (38º570N, 106º590W; ca 2,895m above sea level). Yellow-bellied marmots are hibernating, harem polygynous, facultatively social ground-dwelling squirrels with matrilineal colony structures. This population is active for around five months annually (mid-April to mid-September). Mating soon after emergence from hibernation, new pups emerge, and yearlings disperse in late-June to early-July. Annually, most males and about half of females disperse as yearlings, typically resulting in movement out of our study area (Armitage 2014). Marmots were studied annually at seven colony sites spread across 5 km at the bottom of the valley. Colonies are grouped into higher and lower elevation sites (four are at higher and three are at lower elevation sites). Higher elevation sites are approximately 166 m higher and experience harsher weather conditions (Van Vuren & Armitage 1991; Blumstein et al. 2006; Maldonado-Chaparro et al. 2015).

Throughout the active season, marmots were repeatedly live trapped, and their social behaviour systematically quantified from 2003 to 2021. Recapture rate is above 86% in all colonies for all sex and age classes considered (Ozgul et al 2006). In addition to unique ear tags, all individuals were marked with unique non-toxic dye marks on their dorsal pelage to allow for accurate identification. Marmots were weighed when trapped and these data were used to predict 1 June and 15 August body mass (to estimate early and late season body condition) via a best linear unbiased predictions (BLUPs) model (Ozgul et al. 2010; Kroeger et al. 2018). June mass reflects the energy trade-off between left over energy from hibernation and available energy for spring reproduction whereas August mass reflects gain in fat mass during the active season and predicts overwinter survival (Ozgul et al. 2010; Armitage 2014; Kroeger et al. 2018). Data used in our BLUPs consisted of 25,979 observations across 4,330 individuals and 58 years with a mean of 5.99 observations per individual (range: 1.0–107.0; Median = 4.0), facilitating the accuracy and reliability of the BLUPs (Martin & Pelletier 2011; Dingemanse et al. 2019; Philson et al. 2022).

### (b) Social Networks

Detailed behavioural observational methodology and the ethogram are outlined in Blumstein et al. (2009). For social interactions, the initiator, recipient, location, time, and type of each interaction were recorded, with most interactions (79%) occurring between identified individuals. The remaining 21% interactions could not be identified because of the interacting individuals’ posture or visual obstructions. We excluded these interactions from our data, which should not significantly influence our estimates of social structure (Silk et al. 2015; Sánchez-Tójar et al. 2018). We only included adults and yearlings because only these cohorts were present in spring, when social interactions were most common, and because pups emerge mid-season and primarily interact with their mother and each other (Nowicki & Armitage 1979). We eliminated transients by excluding individuals observed or trapped fewer than five times in a given year (Wey & Blumstein 2012; Fuong et al. 2015; Yang et al. 2017; Blumstein et al. 2018). Only interactions in April, May, and June (∼2.5-month time frame is when marmots emerge from hibernation/mate to when pups emerge from natal burrows) were used because this is when most social interactions occur and when we have the highest resolution of observation data (the growth of vegetation begins to impair observations as the summer progresses). Lastly, we focused on affiliative interactions (e.g. allogrooming, greeting, play) because they relate to fitness on both the individual and group levels (Wey & Blumstein 2012; Yang et al. 2017; Blumstein et al. 2018; Montero et al. 2020; Philson et al. 2022; Philson & Blumstein 2023a; Philson & Blumstein 2023b) and affiliative interactions comprised 88% of all social interactions.

Marmots share space with a subset of all possible individuals within their colony area. We therefore defined social groups based on space-use overlap per year (two individuals observed using the same burrow or seen/trapped at the same location and time within one-day intervals). Simple-ratio pairwise association indices based on colony space-use overlap were (Cairns & Schwager 1987) calculated with SOCPROG (version 2.9; Whitehead 2009) and run through the random walk algorithm Map Equation (Csardi & Nepusz, 2006; Rosvall & Bergstrom, 2008; Rosvall et al., 2009) to assemble association indices and identify social groups (network isolates within an association index). While Map Equation assigns each individual to only one social group (per year in our case), this can exclude key social connections, such as those with adult males. Because adult males often mate with females from multiple matrilines and have important interactions with members of multiple groups, we added adult males to each social group for which they had at least one social interaction with a member of that group to enable more accurate social network measures. However, each year, a male’s network measures were only calculated from their originally assigned group.

From these spatially defined groups, directed and weighted social interaction matrices were constructed from affiliative social interactions for each group each year with the R (version 4.2.0; R Development Core Team 2023) package “igraph” (version 1.4.2; Csardi & Nepusz 2006). These affiliative social interaction matrices consisted of 42,369 social interactions between 1,294 individuals (338 of whom were observed across multiple years). This operationalisation produced 180 social groups with group sizes ranging from 2 to 35 individuals with a mean of 7.65 ± 5.92 (mean ± standard deviation). Individuals had an average of 66.23 ± 90.72 social interactions per year, ranging from 1 to 694. Within social groups, social interactions averaged 447.35 ± 653.18, ranging from 2 to 4,118.

We calculated (using “igraph”) four pairs of analogous individual and group social network measures to quantify the independent contributions of the individual and group phenotypes (Table 1). These analogous network measure pairs quantify four core traits in human and non-human animal social networks (including yellow-bellied marmots) at the two levels of social network organisation: connectivity, closeness, breakability, and clustering (Wasserman & Faust 1994; Krause et al. 2009). Degree (how may social partners an individual has; Wasserman & Faust 1994) and density (proportion of possible social connections that are observed in a group; Opsahl 2009) are paired to quantify social connectivity. Closeness (social distance between all other individuals in the group; Wasserman & Faust 1994) and average path length (mean social distance between all individuals in the group; Opsahl 2009) quantify social closeness. Embeddedness (individual connectedness based on their direct and indirect relationships with their cluster and group; Moody & White 2003) and cut points (number of social links that if broken or lost result in two or more social groups of at least two individuals; Borgatti 2006) quantify social breakability. Local clustering coefficient (proportion of an individual’s direct social partners who are also social partners; i.e., local transitivity; Croft et al. 2008) and transitivity (proportion of possible social triads that are observed in a group; i.e. global transitivity; Wey et al. 2019) quantify social clustering. After scaling (see *Contextual Analysis* below), we flipped the sign for average path length and cut points, so all social network measures’ values correspond to increased sociality. This was done to facilitate interpretation and presentation as the directional slope in Figure 1 can be interpreted as the direction of selection for all measures. We thus refer to inverse average path length and inverse cut points throughout.

**Figure 1.**
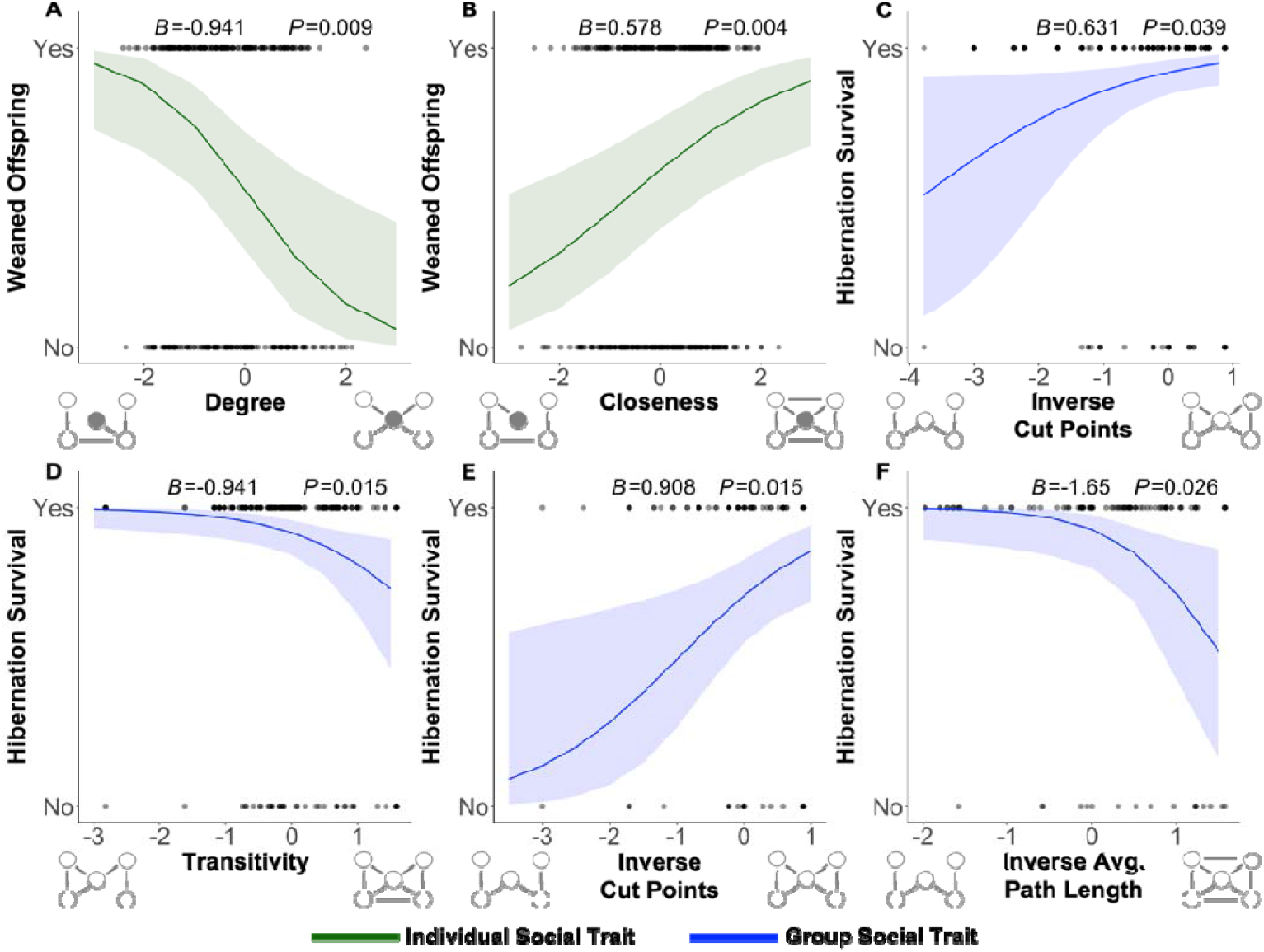
Selection gradients (plotted as marginal effects) for social traits representing the individual and group social phenotypes. For the individual social phenotype (green), connectivity was under negative selection (A; fewer social partners) and closeness was under positive selection (B; less social distance from others) for adult female reproductive success. For the group social phenotype (blue), adult females were under positive selection for breakability (C; more social ties required to break to split into two or more groups) and negative selection for clustering (D; lower proportion of connected transitive loop) for hibernation survival. Group breakability was under positive selection for yearling male (E) hibernation survival. The group closeness trait (mean social distance between all individuals) was under negative selection for adult male hibernation survival (F).

The reliability of the social network measures is facilitated by our regular observations of marmot social groups (mean n observations per individual across years= 28.81, range of each year = 6.79– 75.14) and low rate of unknown individuals involved in social interactions (Silk et al. 2015; Sánchez-Tójar et al. 2018; Davis et al. 2018). Social group size (measured as social network size) is associated with many group-level social network measures (Wasserman & Faust 1994; e.g. density, cut points) which may mediate the strength of selection. Because density, inverse average path length, and transitivity are already “standardised” based on their equations, we standardised inverse cut points by social group size, so all four group-level measures account for social group size. We then fit social group size as a fixed effect in all models to further account for group size.

### (c) Fitness Measures

Summer survival was defined when individuals were seen or trapped after 1 August or in subsequent years and over-winter/hibernation survival was defined as those individuals having survived the summer being seen the following spring or in subsequent years (Philson & Blumstein 2023b). For summer survival, only adults (>2 years old) were included in the analysis because of uncertainty quantifying survival for yearlings because a majority disperse (Armitage 1991). Predation accounts for 98% of summer mortality (Van Vuren 2001) and poor body condition and winter snowpack are primary predictors of hibernation mortality (Van Vuren & Armitage 1991; Armitage et al. 2003). Summer survival was paired with network measures from the current active season and hibernation survival with network measures from the active season before winter.

Only female reproductive success is quantified because male reproductive success mostly depends on dominance, body condition, and tenure length (Armitage 1998; Huang et al. 2011), is difficult to quantify, and the smaller number of males in the population diminishes analysis power. We quantified two attributes of female reproductive success: if a female successfully weaned offspring from the burrow and the number of offspring a mother weaned (if at least one pup was weaned) (Philson & Blumstein 2023a). Behavioural observations and a comprehensive genetic pedigree (Blumstein et al. 2010; Olson & Blumstein 2010) were used to assign offspring to mothers. This method does not account for pups that may have been born in the burrow but died before emergence (all pups are born in the burrow and emerge ∼30 days after birth; Armitage 2014).

### (d) Contextual Analysis

We used contextual analysis, an extension of the Lande-Arnold selection analysis using partial regression to partition selection among levels (in this case, the individual and group social phenotypes; Lande & Arnold 1983; Goodnight et al. 1992). Contextual analysis is generalisable to multiple levels of organisation pending the lowest level included is the level at which fitness variation is being explored. Our contextual analysis differed from classic contextual analysis (Heisler & Damuth 1987; Goodnight et al. 1992; Goodnight & Stevens 1997) since we used emergent group traits instead of using the mean of all individuals within a group.

Contextual analyzes are sensitive to the scale of standardization and should be based on the biological and ecological processes that generate selection in the context of the study system (De Lisle & Svensson 2017; Costello et al. 2023). Thus, we mean-variance standardised individual-level social network measures at the scale of each social group. Because group-level selection inherently operates across groups, we standardised group-level social network measures across all social groups across all 19 years. We further mean-variance standardised group size on the global scale across all social groups across all years. Overall, the model can be expressed as:

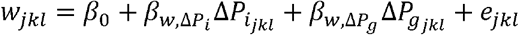

Where *w*_*jkl*_ is the relative fitness of individual *j* in group *k* in year *l, P*_*i*_ are the social traits of an individual (individual social phenotype) and *P*_*g*_ are the social traits of the social group (group social phenotype). 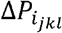 is the deviation of social trait for individual *j* from the mean of its group *k* in year *l*. 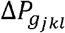 is the deviation of the social group trait for group *k* of individual *j* in year *l* from the overall mean of social group trait across all groups and all years. 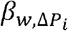 and, 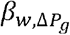 are the among-individual and among-group selection gradients. and *e*_*jkl*_ are the intercept and residual terms respectively (Lande & Arnold 1983; Fisher et al. 2017).

Due to fundamental differences in life history strategies between age classes (adult and yearling) and sexes in this system (Armitage 2014), we fit separate models for each cohort. Each model included the fitness measure as the response variable, the four pairs of analogous social network traits, social group size, and valley location (higher or lower elevation) as fixed effects to account for known environmental and social variation. Individual ID and year were fit as random effects to further account for environmental and demographic variation. Eight models were fit in total and the final models, after correcting fit for multicollinearity, met their respective assumptions. In R (version 4.2.0; R Development Core Team 2023), models were fit with “lme4” (version 1.1-33; Bates et al. 2015) and assumptions were checked with the packages “car” (version 3.1-2; Fox & Monette 1992; Fox & Weisberg 2019) and “DHARMa” (version 0.4.6; Hartig 2012).

The set of models (adult females only) for both measures of reproductive success additionally included June body mass as a fixed effect given body condition’s importance for reproductive success in this system (Armitage 2014); neither model had multicollinearity issues. The model for whether or not a female weaned a litter was fit with a binomial distribution with a bobyqa optimizer of 10,000 function evaluations and had 363 observations across 157 unique individuals. The model for the number of offspring (if a litter was weaned) was fit with a Poisson distribution and had 191 observations across 98 unique individuals (Supplementary Table 1).

The four sets of models (one for each age-sex cohort) for hibernation survival additionally included August body mass as a fixed effect (Armitage 2014). These models were fit with a binomial distribution and a bobyqa optimizer of 10,000 function evaluations. Multicollinearity was an issue between the network traits and thus the degree-density analogous pair was removed from all four models. With 19 years of data, the yearling male and female models had 119 and 209 observations, respectively (Supplementary Table 2). The adult male model had 109 observations across 58 unique individuals and the adult female model had 324 observations across 134 unique individuals across 19 years (Supplementary Table 3).

The two models for summer survival (male and female adults only) additionally included June body mass and a predator index as fixed effects (Armitage 2014). The predation index was calculated by dividing the number of predators seen in a colony by the time spent observing that colony for that year (Blumstein et al. 2023). These models were fit with a binomial distribution and a bobyqa optimizer of 10,000 function evaluations. Both of these models had VIF > 5 for density, thus we removed the degree-density analogous pair and ran the models again. The summer model included 19 years of data with 138 observations across 80 unique individuals for males and for females had 363 observations across 157 unique individuals (Supplementary Table 4).

Because of the standardization approach we used as part of this contextual analysis, model estimates represent the magnitude of the selection gradient. To account for potential non-linear selection, we re-fit all models with a quadratic transformation for both the individual and group social network measures as fixed effects in each model. However, because these non-linear variables were not statically significant in any model, we chose to report the models without these non-linear variables for model fit, statistical simplicity, and interpretability. We also controlled for multiple comparisons by calculating False Discover Rate (FDR) adjusted *P*-values (Shaffer 1995; Verhoeven et al., 2005; Pike, 2011) based on eight comparisons for the eight models, which still showed multilevel selection occurred (despite losing statistical significance for some group measures after the FDR adjustment; Supplementary Table 5).

In summary, our contextual analysis (1) accounts for the inherent non-independence of the individual and group social phenotypes by partitioning among the two phenotypic levels relative to each other, and (2) controls for other biologically relevant variables that, if excluded, could otherwise cause spurious associations. If we find significant evidence of selection, it is within the context of the biology of this system, not in isolation. Further, given social groups are largely comprised of kin (hence the matrilineal nature of marmot society), kin selection (which seeks to explain optimality while multilevel selection does not) is not relevant for our question and group size (as groups are mostly kin) in part controls for the number of kin (Goodnight 2013). Our contextual analysis permits the quantification of independent contributions to individual fitness at different social phenotypic levels, and between age-sex cohorts, while accounting for natural variation in environmental and demographic factors across a nearly two-decade dataset of a wild, free-living mammal.

## 3. Results and Discussion

### (a) Context-Dependent Multilevel Selection

Our results revealed quantitative differences in the presence, strength, and direction of multilevel selection for individual and group social phenotypes dependent on sex, age, and fitness context (Figure 1).

Individual connectivity was under negative selection (i.e., fewer direct social partners) and individual closeness was under positive selection (i.e., socially closer to all others in the group, directly and indirectly) for reproductive success in adult females (Figure 1A-B; Supplementary Table 1). This could represent an evolutionary balance in this system between the time and energetic costs of social relationships for reproductive success (Wey and Blumstein 2012), while still maintaining closer social distance to others to maximize anti-predator benefits (e.g., hearing alarm calls), allowing for more time and resources to be devoted to reproduction (Montero et al. 2020). Given each species’ time budget and selective pressures are unique, what individual social traits are under selection, and in what way, will be context dependent based on the biology of that system. For example, in more gregarious species, more aligned selection across individual social traits may be observed.

Group clustering was under negative selection (i.e., there were fewer connected transitive loops) for adult female hibernation survival (Figure 1D; Supplementary Table 3), showing selection for adult females residing in less socially connected and clustered social groups. Group closeness was also under negative selection (i.e., larger social distance between all individuals) for adult male hibernation survival (Figure 1F; Supplementary Table 3). This shows selection for adult males residing in groups where all individuals are more socially distant. Yellow-bellied marmots are facultatively social, meaning compared to more gregarious species marmots are more socially flexible and sometimes solitary (Armitage 2014). Thus, marmots may not benefit from residing in more connected and social groups, as supported by this negative selection for social structures that may facilitate increased rates of social interactions. However, group breakability was under positive selection (i.e., more social ties required to break a group into two or more separate groups) for both adult female (Figure 1C) and yearling male (Figure 1E) hibernation survival (Supplementary Table 2 and 3). That is, selection favors individuals residing in more structurally cohesive and less fragmentable social groups because these groups may facilitate synchrony in hibernation onset (Arnold 1993) and because the breaking of social groups may reduce the number of individuals engaging in social hibernation, reducing subsequent thermoregulatory benefits and increasing the risk of death in hibernation (as seen in the alpine marmot (*Marmota marmota*); Arnold 1993). Collectively, this suggests group structures that facilitate phenological timing and thermoregulatory benefits are under positive selection, while structures that foster increased rates of interaction are under negative selection. The particular type of group social structure matters.

Compared to exploring the nested levels of social organisation independently, which may lead to an inaccurate estimation of selection, this multilevel selection analysis revealed a more comprehensive picture of selection for sociality in this system. Namely, only the two group-level breakability (i.e., inverse cut points) results were found previously (Philson and Blumstein 2023b). This difference could be attributable to multiple factors, such as different sized datasets and statistical approaches across the studies; the individual social phenotype and reproductive success analysis was conducted over a decade ago (Wey and Blumstein 2012). Now with a 19-year dataset and a standardised statistical approach, we have a more accurate estimate of selection for sociality in this system. While exploring the levels of sociality independently provided a strong background to ask informed questions about the importance of multilevel selection for social traits, this multilevel selection approach, partitioning selection among the nested levels of organisation, relevels a more informed and representative view of natural selection in this system, facilitating more accurate evolutionary predictions about sociality in this system.

Results somewhat align with our *a priori* predictions of negative selection for less connected, close, and clustered social traits at both phenotypic levels and positive selection for less breakable traits at both phenotypic levels. However, individual closeness was under positive selection, contrary to our prediction. Neither social phenotypic level was under selection for summer survival (Supplementary Table 4) or in female yearlings (Supplementary Table 2), also contrary to our *a priori* predictions. Because the two social phenotypic levels are scaled, model estimates represent the magnitude of the selection gradient. Across all traits, age classes, and both sexes, the individual social phenotype had a mean selection gradient of 0.76 (0.26 SE) and the group social phenotype a mean of 1.03 (0.43 SE). This suggests that selection is just as strong, if not stronger for the group social phenotype, contrary to our *a priori* prediction and to the classic criticism of multilevel selection. Collectively, these results support the importance of a multilevel selection framework to better understand the evolution and maintenance of sociality.

### (b) Antagonistic Multilevel Selection on Sociality

Not only do these results suggest multilevel selection may be occurring in the wild for individual social behaviour and group social structure, but that there is antagonistic multilevel selection *within* each of the two phenotypic levels and *between* the two phenotypic levels (Weinig et al., 2007; Eldakar et al., 2010; Costello et al. 2023). For adult females, there is both positive and negative selection within the individual (between connectivity and closeness) and within the group social phenotype (between closeness, breakability, and clustering). There is also antagonistic selection between the two social phenotypes for adult females: positive selection for more social individuals potentially being counteracted by selection for less social groups, and *vice versa*. Antagonistic multilevel selection *within* and *between* social phenotypes may flatten or constrain the overall selection pressure (i.e. leading to stabilizing selection) for both social phenotypes (Weinig et al., 2007; Goodnight 2013), which could explain why sociality is not as highly heritable (*h*^*2*^ = 0.11; Lea et al. 2010) or ubiquitously beneficial in this system as it is in other, more gregarious systems (Snyder-Mackler et al. 2020), such as humans (*Homo sapiens*; *h*^*2*^ = 0.47; Fowler et al. 2009) and rhesus macaques (*Macaca mulatta*; *h*^*2*^ = 0.84; Brent et al. 2013).

In a captive population of forked fungus beetles (*Bolitotherus cornutus*) with a fixed social group size, Costello et al. (2023) showed positive selection for more connected individual social phenotypes in males and negative selection for more connected group social phenotypes in females within the context of reproductive success, suggesting sexually antagonistic multilevel selection (as we also observe, but across different fitness contexts). This study quantified individual and group social phenotypes with a similar analogous social network trait and contextual analysis approach as we do here, but without quantifying the independent contributions of selection for multiple analogous pairs in the same model. Combined with our results, this shows multilevel selection for social phenotypes is present in *Coleoptera* and *Rodentia*. This evidence of multilevel selection in the wild and across species provides further impetus for experimental work to manipulate the composition of individual and group social phenotypes to further disentangle these nested structures, and to nail down causality and eliminate unmeasured variables (Bijma & Wade 2008). For example, future work should leverage wild populations to identify, and experimental populations to confirm, the presence of multilevel selection and the factors that may mediate the strength and context of multilevel selection. Factors such as relatedness, phenotypic plasticity, individual and group age, group composition and sex ratios, and environmental factors (such as resource distribution) could increase or decrease the strength and direction of multilevel selection in other species.

### (c) Multilevel Adaptation in Response to Multilevel Selection

Because the contextual analysis regressed individual fitness against the individual and group social phenotypes (i.e., partitioning selection among the two levels relative to each other), our results of selection at both levels adds evolutionary support to the distinct and discrete nature of individual behaviour and group social structure despite being nested levels of biological organisation. (Okasha 2005; Royle et al. 2012; Goodnight, 2013; Gardner, 2015; Solomon-Lane et al. 2015; Cantor et al. 2021; Philson et al. 2022; Costello et al 2023; Philson & Blumstein 2023a; Philson & Blumstein 2023b). Given we now show selection at multiple phenotypic levels in addition to heritability (Lea et al. 2010) for social behaviour in this system, this allows for the potential of an evolutionary response. Evidence for multilevel adaptation in response to multilevel selection would require demonstration of intergenerational changes in the units of replication that underlie social behaviours (Lande & Arnold 1983). For example, the gene *CG14109* influences betweenness centrality (Rooke et al. 2024), a key property of social networks that indicates an individual’s importance in facilitating disease spread, communication, and cohesion within the network, in fruit flies (*Drosophila melanogaster*). Future work should leverage genomic tools to link individual and group phenotypes with candidate genes, like *CG14109*, and to track intergenerational changes in the units of replication that underlie social behaviours.

Given sociality inherently involves interactions between varying phenotypes, indirect genetic effects likely play a role in the evolution of sociality (Bijma & Wade 2008; Radersma 2021; McLean et al. 2023). Further, in these nested levels of biological organisation, selection at one level may accelerate or constrain evolution at another level (Bijma & Wade, 2008), which is further complicated by the antagonistic multilevel selection within and between social phenotypes we show here. The next steps to better understanding the evolution of societies across species should identify the genetic architecture and correlations underpinning multilevel selection, incorporating both direct and indirect genetic effects in a unified empirical framework (Radersma 2021).

### (d) Group Living and Social Phenotypes

Interestingly we found no selection for either social phenotype in the context of summer survival. Predation accounts for 98% of summer mortality in this system (Van Vuren 2001) and the evolutionary origins of yellow-bellied marmot group living partly involve predator avoidance (Armitage 2014). This suggests selection driving group living can be, and is in some cases, different than the subsequent selection pressures driving how social individuals are and the structure of the social group. Again, work in more gregarious species where the evolutionary origins of group living may be attributable to grooming, heat retention, or resource acquisition could find that the selection for group living and the individual and group social phenotypes are more aligned.

A common argument against group selection (and thus multilevel selection) used since the 1960s focuses on differences in individual lifespan versus group duration (Maynard Smith 1964; Williams 1966; Wade 1977): where individual lifespans are shorter than group duration, selection proceeds faster at the level of the individual. However, in yellow-bellied marmots, dispersal, mortality, and births ensure that groups restructure annually. Individuals surviving to adulthood have a mean lifespan of 4.07 years and are thus members of multiple social groups in their lifetime (Armitage 2014). In this population, individual lifespan is longer than group turnover and thus this argument does not apply. Indeed, the same logic would predict stronger selection for the group phenotype, which we found.

## 4. Conclusion

By leveraging a 19-year dataset from a wild, free-living mammal and contextual analysis to partition selection relatively among two nested levels of sociality, we provide evidence suggestive of multilevel selection as a potentially important and overlooked force in the evolution of sociality. While experimental tests are required to formalize causality and eliminate unmeasured variables affecting fitness and the social phenotypes beyond what our models controlled for, our detection of multilevel selection in the wild provides the impetus to develop these multilevel selection experiments across systems. Considering levels of sociality independently may be reductive and lead to Type I or Type II errors; when we don’t incorporate a multilevel selection perspective, we may incorrectly attribute natural selection to the levels of organisation not actually under direct natural selection. Instead, sociality can be quantified as a nested, multi-dimensional phenotype of which different levels can be selected on and change with age and sex. Compared to exploring levels of sociality independently and given contextual analysis partitions selection among phenotypic levels relative to each other, multilevel selection may more comprehensively represent how natural selection acts on the social interactions and structures that arise from group living. We provide empirical evidence to spark further study in this classic debate in biology.

## Ethics

This study uses a long-term dataset. Marmots were originally studied under the research protocol ARC 2001-191-01 (approved by the UCLA Animal Care Committee on 13 May 2002 and renewed annually), protocols approved by the Rocky Mountain Biological Laboratory, and trapped under permits issued annually by the Colorado Department of Parks and Wildlife (TR-917).

## Supplementary Information

**Supplementary Table 1.**
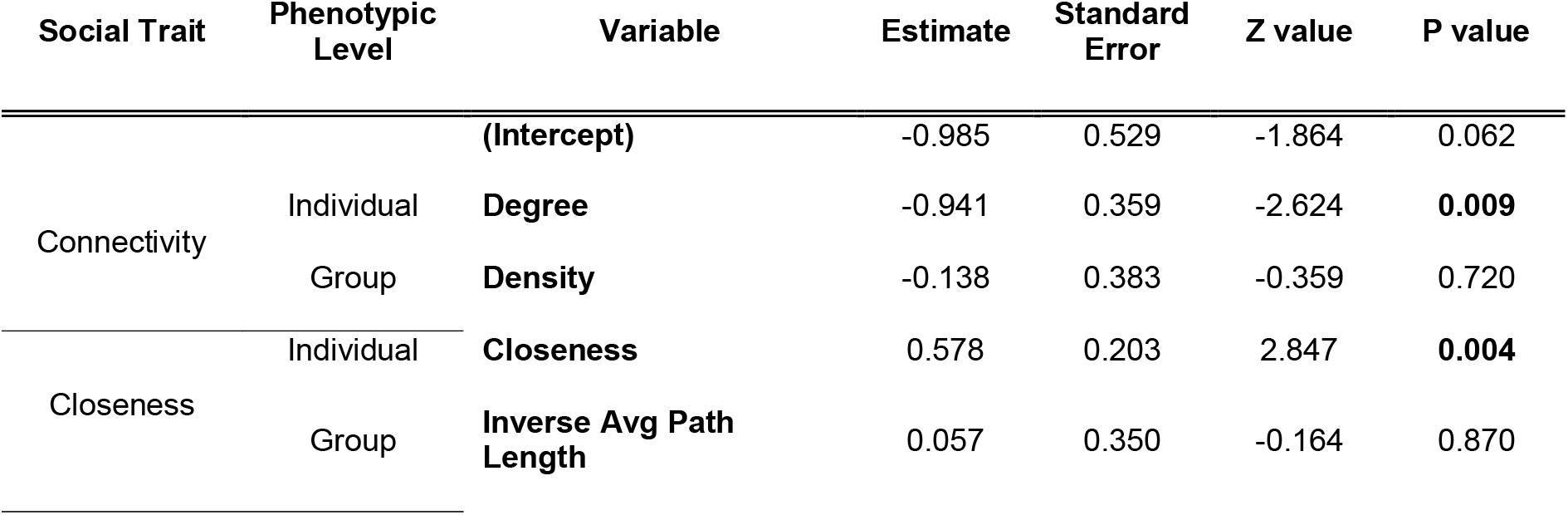
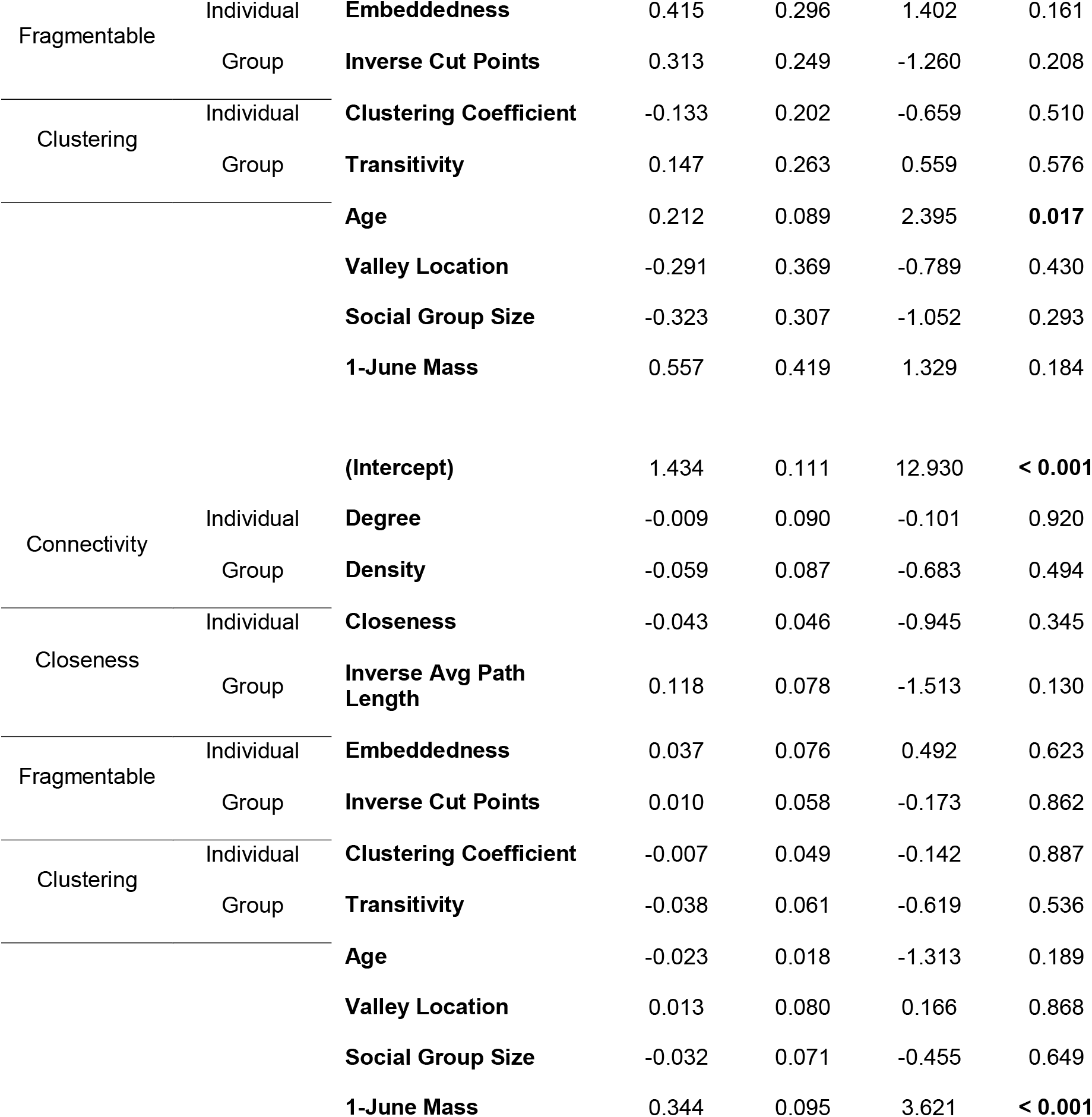
Estimates, standard error, Z value, and P value from the two contextual analyzes for adult female reproductive success: if at least one offspring was successfully weaned from the burrow (top) and the number of offspring a mother weaned (if at least one pup was weaned; bottom).

**Supplementary Table 2.**
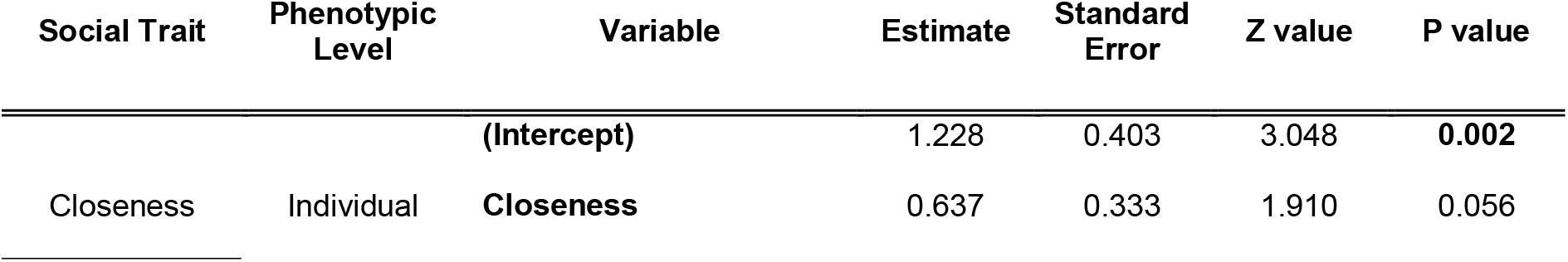
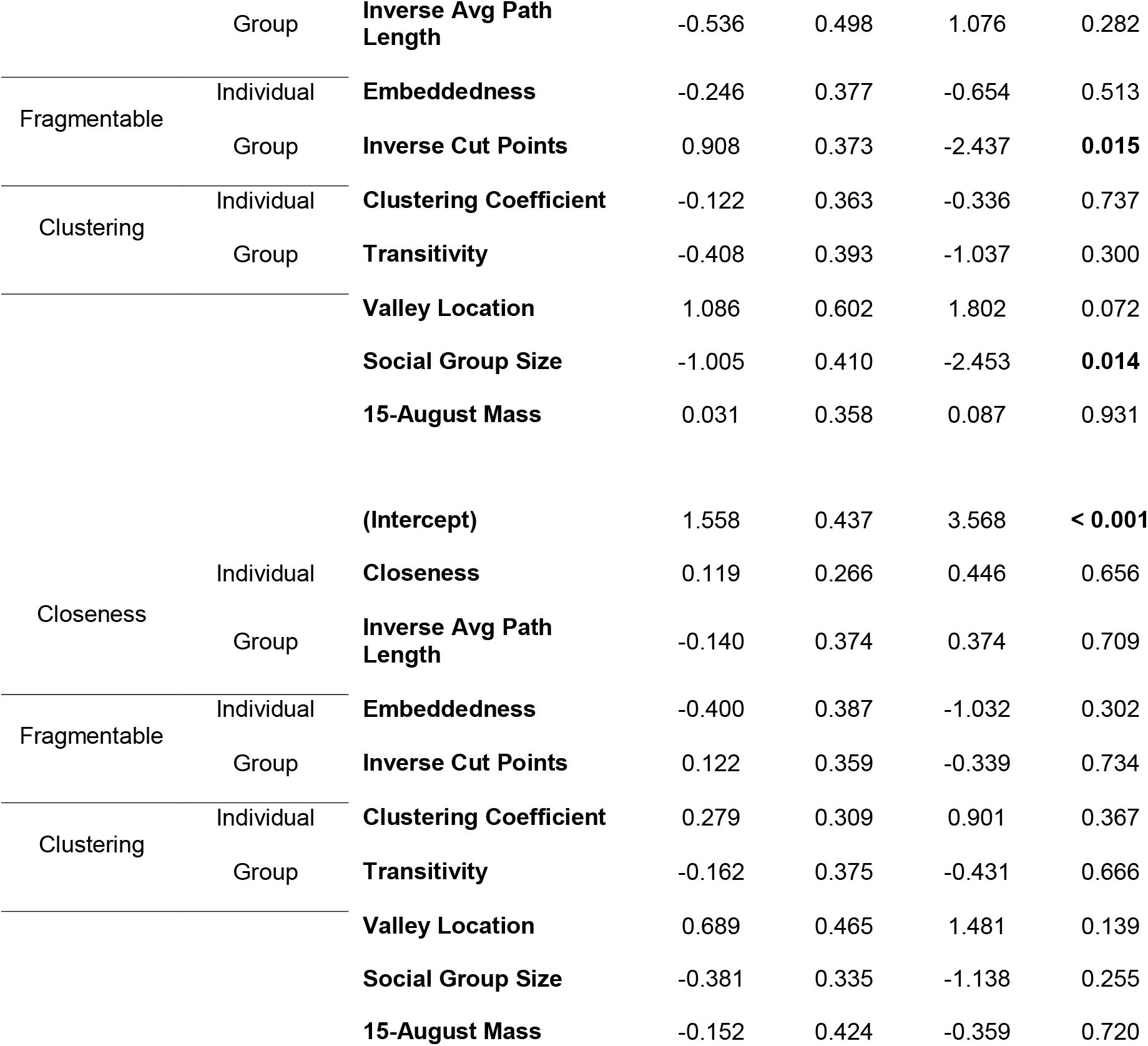
Estimates, standard error, Z value, and P value from the two contextual analyzes for yearling male (top) and yearling female (bottom) hibernation survival.

**Supplementary Table 3.** Estimates, standard error, Z value, and P value from the two contextual analyzes for adult male (top) and adult female (bottom) hibernation survival.

| **Social Trait** | **Phenotypic Level** | **Variable** | **Estimate** | **Standard Error** | **Z value** | **P value** |
| --- | --- | --- | --- | --- | --- | --- |
|  |  | **(Intercept)** | 1.027 | 0.865 | 1.188 | 0.235 |
| Closeness | Individual | **Closeness** | -0.327 | 0.389 | -0.840 | 0.401 |
|  | Group | **Inverse Avg Path Length** | -1.648 | 0.739 | 2.228 | **0.026** |
| Fragmentable | Individual | **Embeddedness** | 0.716 | 0.480 | 1.491 | 0.136 |
|  | Group | **Inverse Cut Points** | -0.074 | 0.485 | 0.152 | 0.879 |
| Clustering | Individual | **Clustering Coefficient** | 0.209 | 0.356 | 0.588 | 0.556 |
|  | Group | **Transitivity** | 0.060 | 0.493 | 0.121 | 0.904 |
|  |  | **Valley Location** | 0.003 | 0.690 | 0.005 | 0.996 |
|  |  | **Social Group Size** | -1.378 | 0.682 | -2.022 | **0.043** |
|  |  | **15-August Mass** | 0.747 | 0.390 | 1.918 | 0.055 |
|  |  | **(Intercept)** | 2.498 | 0.418 | 5.971 | **< 0.001** |
| Closeness | Individual | **Closeness** | -0.388 | 0.244 | -1.591 | 0.112 |
|  | Group | **Inverse Avg Path Length** | -0.001 | 0.372 | 0.001 | 0.999 |
| Fragmentable | Individual | **Embeddedness** | 0.316 | 0.230 | 1.371 | 0.170 |
|  | Group | **Inverse Cut Points** | 0.631 | 0.306 | -2.063 | **0.039** |
| Clustering | Individual | **Clustering Coefficient** | -0.079 | 0.219 | -0.361 | 0.718 |
|  | Group | **Transitivity** | -0.941 | 0.385 | -2.444 | **0.015** |
|  |  | **Valley Location** | -0.168 | 0.420 | -0.400 | 0.689 |
|  |  | **Social Group Size** | -0.647 | 0.358 | -1.806 | 0.071 |
|  |  | **15-August Mass** | 0.144 | 0.303 | 0.477 | 0.634 |

**Supplementary Table 4.** Estimates, standard error, Z value, and P value from the two contextual analyzes for adult male (top) and adult female (bottom) summer survival.

| **Social Trait** | **Phenotypic Level** | **Variable** | **Estimate** | **Standard Error** | **Z value** | **P value** |
| --- | --- | --- | --- | --- | --- | --- |
|  |  | **(Intercept)** | 0.840 | 0.685 | 1.226 | 0.220 |
| Closeness | Individual | **Closeness** | 0.333 | 0.299 | 1.114 | 0.266 |
|  | Group | **Inverse Avg Path Length** | 0.618 | 0.434 | -1.424 | 0.154 |
| Fragmentable | Individual | **Embeddedness** | 0.272 | 0.318 | 0.854 | 0.393 |
|  | Group | **Inverse Cut Points** | 0.022 | 0.318 | -0.070 | 0.944 |
| Clustering | Individual | **Clustering Coefficient** | -0.072 | 0.277 | -0.261 | 0.794 |
|  | Group | **Transitivity** | -0.026 | 0.371 | -0.070 | 0.944 |
|  |  | **Valley Location** | 0.175 | 0.531 | 0.329 | 0.742 |
|  |  | **Social Group Size** | -0.135 | 0.434 | -0.310 | 0.756 |
|  |  | **1 June Mass** | 0.551 | 0.318 | 1.733 | 0.083 |
|  |  | **Predator Index** | -0.339 | 0.528 | -0.642 | 0.521 |
|  |  | **(Intercept)** | 2.108 | 0.484 | 4.357 | **< 0.001** |
| Closeness | Individual | **Closeness** | -0.112 | 0.230 | -0.488 | 0.625 |
|  | Group | **Inverse Avg. Path Length** | -0.323 | 0.357 | 0.905 | 0.366 |
| Fragmentable | Individual | **Embeddedness** | 0.412 | 0.215 | 1.920 | 0.055 |
|  | Group | **Inverse Cut Points** | 0.156 | 0.264 | -0.588 | 0.556 |
| Clustering | Individual | **Clustering Coefficient** | -0.187 | 0.208 | -0.900 | 0.368 |
|  | Group | **Transitivity** | 0.305 | 0.262 | 1.162 | 0.245 |
|  |  | **Valley Location** | -0.567 | 0.447 | -1.268 | 0.205 |
|  |  | **Social Group Size** | -0.344 | 0.365 | -0.943 | 0.346 |
|  |  | **1 June Mass** | 1.156 | 0.382 | 3.027 | **0.002** |
|  |  | **Predator Index** | 0.095 | 0.401 | 0.238 | 0.812 |

**Supplementary Table 5.** False Discover Rate (FDR) adjusted P-values (for eight comparisons due to running eight models) for the six statistically significant social network measures.

| **Fitness Measure** | **Network Measure** | **Estimate** | **Standard Error** | **Z value** | **P value** | **FDR Adjusted P value** |
| --- | --- | --- | --- | --- | --- | --- |
| If Offspring Was Weaned | **Closeness** | 0.578 | 0.203 | 2.847 | 0.004 | **0.035** |
| If Offspring Was Weaned | **Degree** | -0.941 | 0.359 | -2.624 | 0.009 | **0.035** |
| Hibernation Survival<br>(Adult Female) | **Transitivity** | -0.941 | 0.385 | -2.444 | 0.015 | **0.039** |
| Hibernation Survival<br>(Yearling Male) | **Inverse Cut Points** | 0.908 | 0.373 | -2.437 | 0.015 | **0.030** |
| Hibernation Survival<br>(Adult Male) | **Inverse Avg. Path Length** | -1.648 | 0.739 | 2.228 | 0.026 | **0.041** |
| Hibernation<br>Survival<br>(Adult Female) | **Inverse Cut**<br>**Points** | 0.631 | 0.306 | -2.063 | 0.039 | 0.052 |

